# Assessing white matter plasticity in a randomized controlled trial of early literacy training in preschoolers

**DOI:** 10.1101/2024.08.16.608210

**Authors:** Sendy Caffarra, Iliana I. Karipidis, John Kruper, Emily Kubota, Adam Richie-Halford, Megumi Takada, Ariel Rokem, Jason D. Yeatman

**Affiliations:** Department of Biomedical, Metabolic and Neural Sciences, University of Modena and Reggio Emilia, Modena, Italy; Division of Developmental-Behavioral Pediatrics, Stanford School of Medicine, Stanford, CA, USA; Department of Child and Adolescent Psychiatry and Psychotherapy, University Hospital of Psychiatry Zurich, Zurich, Switzerland; Neuroscience Center Zurich, University of Zurich and ETH, Zurich, Switzerland; Department of Psychiatry and Behavioral Sciences, Stanford School of Medicine, Stanford, CA, USA; Department of Psychology, University of Washington, Seattle, WA, USA; eScience Institute, University of Washington, Seattle, WA, USA; Graduate School of Education, Stanford University, Stanford, CA, USA

**Author notes:** Corresponding author: Sendy Caffarra (SC).

## Abstract

Reading is a cognitive skill that requires our brain to go through a myriad of changes during learning. While many studies have described how reading acquisition shapes children’s brain function, less is known about the impact of reading on brain structure. Here we examined short-term causal effects of reading training on preschoolers’ behavior and white matter structure. Forty-eight English-speaking preschoolers (4y10m to 6y2m) participated in a randomized controlled trial where they were randomly assigned to two training programs: the Letter training program was focused on key skills for reading (e.g., decoding and letter knowledge), while the Language training program strengthened oral language comprehension skills without exposure to text. Longitudinal behavioral data showed that only the Letter Training group increased letter knowledge and decoding skills after the two-week training. Diffusion MRI measures (FA and MD) of eighteen white matter pathways (including the left arcuate and the left inferior longitudinal fasciculus) did not reveal any statistically significant changes for either group despite high degrees of scan-rescan reliability across sessions. These findings suggest that a two-week reading training program can cause changes in preschoolers’ letter knowledge and decoding abilities, without being accompanied by measurable changes in the diffusion properties of the major white matter pathways of the reading network. We conclude highlighting possible constraints (i.e., age, training onset and duration, cognitive profile) to reading-related white matter plasticity.

## Introduction

Reading is a complex cognitive skill that has an impact on brain structure and function. Learning to decode written language not only changes the way the brain functions [1,2], but is also associated with changes in the structural properties of white matter pathways [3–5]. However, it is still unclear under which conditions (e.g., quantity and quality of the training, developmental stage), and at what time scale experience-dependent structural changes emerge. In addition, the relationship between learning-driven changes in reading behavior and brain plasticity is still underspecified. To deepen our understanding of the short-term effects of the initial phase of reading acquisition on brain structure, we ran a diffusion magnetic resonance study (dMRI) that used a randomized controlled trial in preschoolers to test how training in letter-speech sound knowledge affects behavior, white matter structure, and their relationship over the course of two weeks.

A growing body of research reports a relationship between reading experience and structural properties of white matter pathways [6]. Among the white matter tracts of the reading circuitry there are the left arcuate (AF), connecting language areas in frontal and temporal lobes [7], and the left inferior longitudinal fasciculus (ILF), which connects the anterior temporal cortex to more posterior ventral-occipital areas (including the visual word form area, [8]). Diffusion properties of left AF and ILF, such as fractional anisotropy (FA) and mean diffusivity (MD), have been related to reading performances in a single time point (concurrent or prior to the dMRI acquisition, [9–11]), as well as over a series of longitudinal observations [4,5,12,13]. Recent studies have started to highlight the presence of a dynamic relationship between the longitudinal trajectories of white matter structure and changes of reading performance over time [14,15]. For instance, studies focused on long-term reading-related structural plasticity reported that typically developing children show increased FA and/or decreased MD in reading white matter pathways (e.g., left AF and left ILF) as reading scores improve. These structural changes are evident over one to four years of formal reading instruction [11,13,16–18]. Moreover, the rate of FA change in the left AF relates to the rate of change in reading performances over a period of five years of reading instruction [15].

Longitudinal studies focused on a shorter period of reading training have led to mixed findings. Table 1 summarizes the available longitudinal research on reading-dependent changes of white matter diffusion properties. Unfortunately, the picture provided by these studies is only partial since most research so far has been focused on reading interventions for children with (or at-risk of) reading disorders and its short-term effects on white matter properties.

**Table 1.**
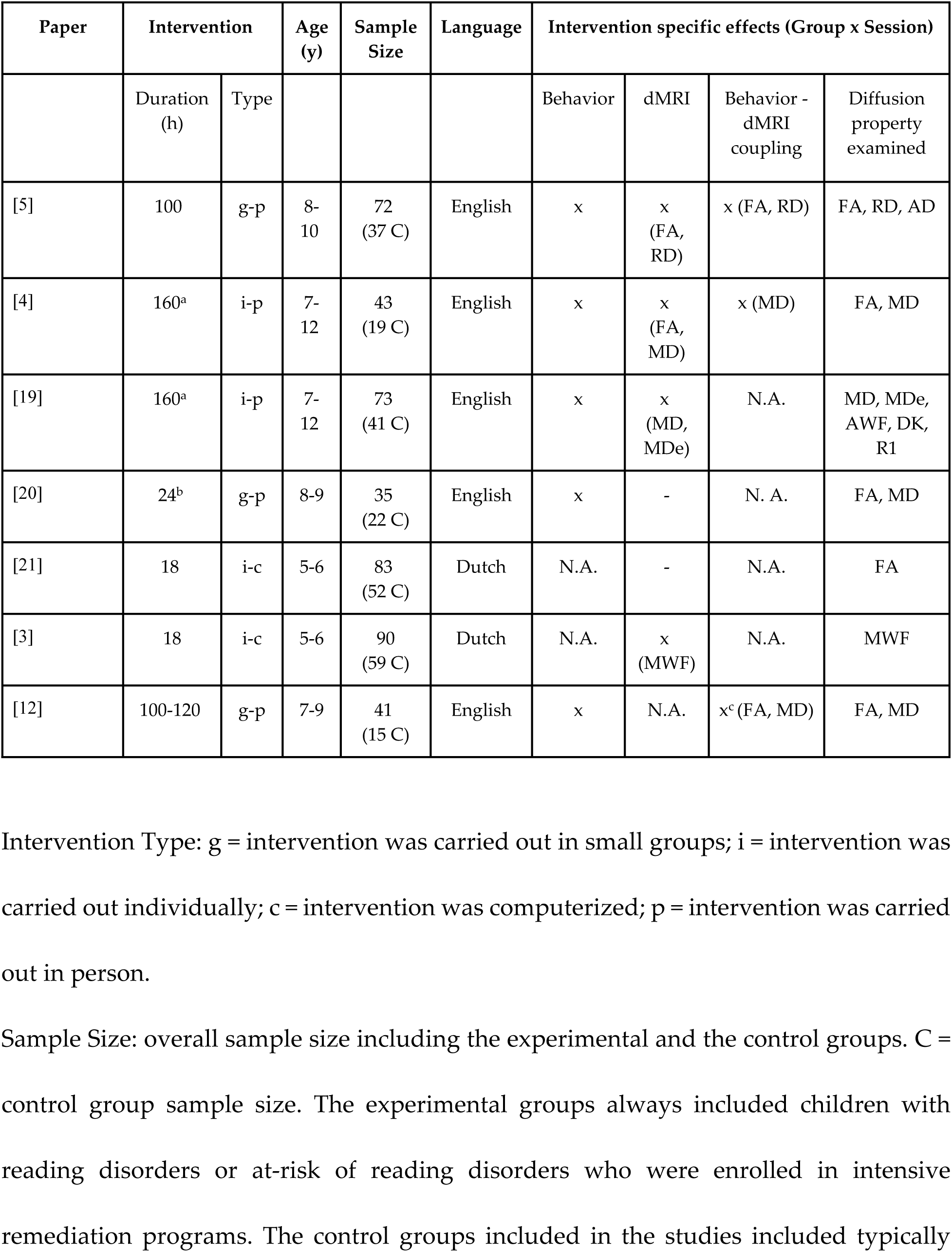
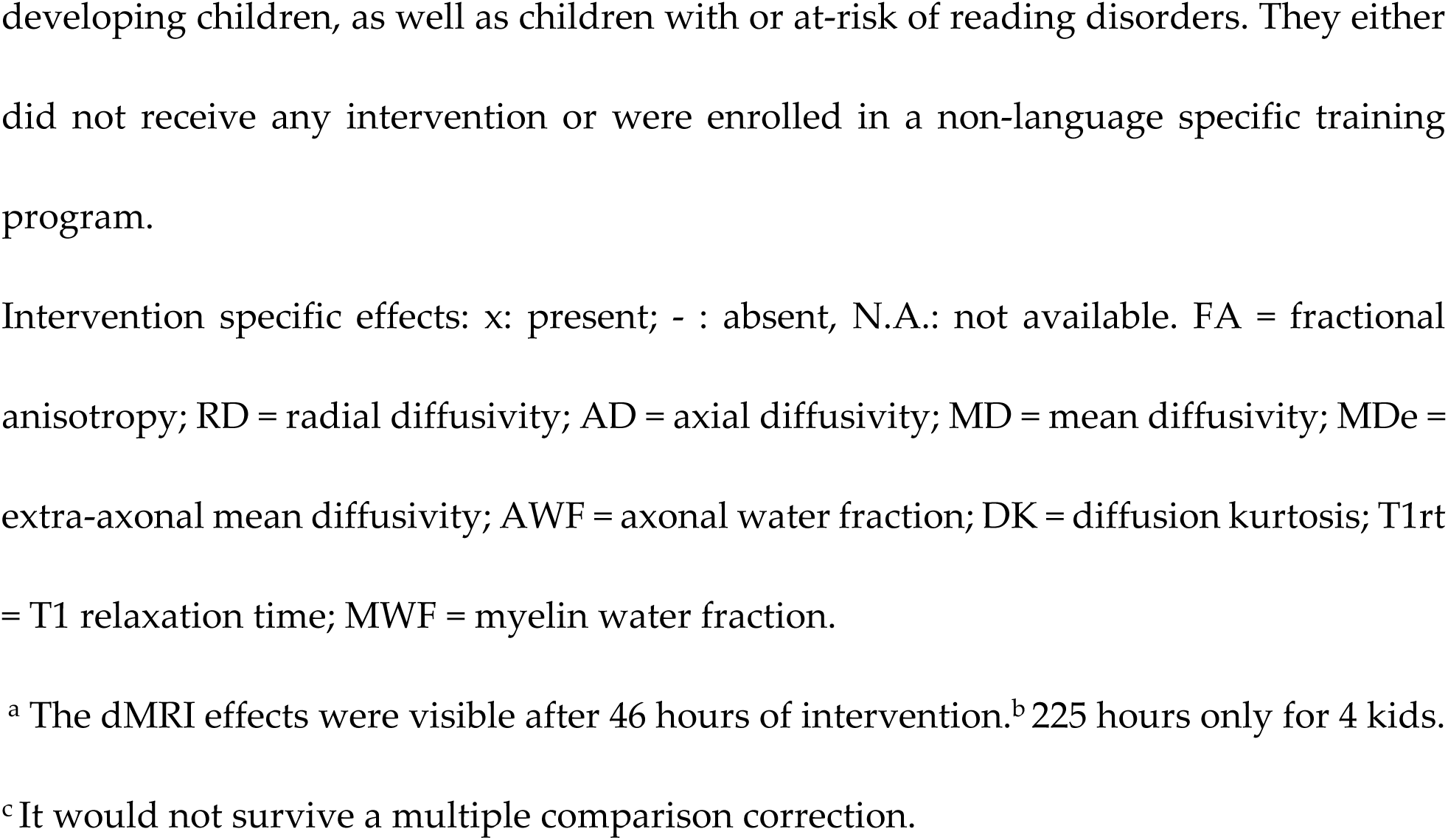
Studies on white matter changes due to short-term reading intervention programs.

| Paper | Intervention |  | Age (y) | Sample Size | Language | Intervention specific effects (Group x Session) |  |  |  |
| --- | --- | --- | --- | --- | --- | --- | --- | --- | --- |
|  | Duration (h) | Type |  |  |  | Behavior | dMRI | Behavior - dMRI coupling | Diffusion property examined |
| [5] | 100 | g-p | 8-10 | 72 (37 C) | English | x | x (FA, RD) | x (FA, RD) | FA, RD, AD |
| [4] | 160<sup>a</sup> | i-p | 7-12 | 43 (19 C) | English | x | x (FA, MD) | x (MD) | FA, MD |
| [19] | 160<sup>a</sup> | i-p | 7-12 | 73 (41 C) | English | x | x (MD, MDe) | N.A. | MD, MDe, AWF, DK, R1 |
| [20] | 24<sup>b</sup> | g-p | 8-9 | 35 (22 C) | English | x | - | N. A. | FA, MD |
| [21] | 18 | i-c | 5-6 | 83 (52 C) | Dutch | N.A. | - | N.A. | FA |
| [3] | 18 | i-c | 5-6 | 90 (59 C) | Dutch | N.A. | x (MWF) | N.A. | MWF |
| [12] | 100-120 | g-p | 7-9 | 41 (15 C) | English | x | N.A. | x<sup>c</sup> (FA, MD) | FA, MD |

Intervention Type: g = intervention was carried out in small groups; i = intervention was
carried out individually; c = intervention was computerized; p = intervention was carried
out in person.
Sample Size: overall sample size including the experimental and the control groups. C =
control group sample size. The experimental groups always included children with
reading disorders or at-risk of reading disorders who were enrolled in intensive
remediation programs. The control groups included in the studies included typically
developing children, as well as children with or at-risk of reading disorders. They either did not receive any intervention or were enrolled in a non-language specific training program.
Intervention specific effects: x: present; - : absent, N.A.: not available. FA = fractional anisotropy; RD = radial diffusivity; AD = axial diffusivity; MD = mean diffusivity; MDe = extra-axonal mean diffusivity; AWF = axonal water fraction; DK = diffusion kurtosis; T1rt = T1 relaxation time; MWF = myelin water fraction.

Although the number of studies conducted on this topic is still low, some preliminary trends can be highlighted given the available findings. First, all studies consistently showed that only children who received reading intervention improved their reading performance, confirming the efficacy of short-term reading training programs [4,5,12,19,20]. Second, these intervention-specific behavioral changes were not always accompanied by fast learning-driven white matter changes, suggesting that behavioral and brain structure changes do not necessarily co-occur or this co-occurrence might be specific to a subset of white matter diffusion properties [3–5,19–21]. Third, there is scarce and mixed evidence on how effects generalize across different age ranges and types of training, pointing to the need for additional research on short-term coupling between reading behavior and white matter plasticity [4,5,12]. Finally, all studies listed above focused on remediation programs and large effects were seen with intensive (high dosage) training in late elementary school children [4,12]. Hence, they provide insights on rapid white matter changes that might also reflect compensatory mechanisms of the reading circuitry, or other factors that are unique to older children with dyslexia, rather than solely plasticity due to the experience of learning to read.

The present exploratory study aims to complement the available research on short-term white matter plasticity by focusing on language and literacy training programs in typically developing preschool children. A randomized controlled trial was conducted with English-speaking preschoolers who were enrolled in a two-week program which either trained reading or spoken language skills (Letter and Language program, respectively). Behavioral and dMRI measures were collected before and after the training to test how preschool learning might affect changes in the white matter. To better characterize the quality and consistency of children’s dMRI measures over time, scan-rescan reliability metrics were calculated for each dMRI measure (FA and MD) and white matter tract.

## Materials and Methods

### Participants

Forty-eight English-speaking preschoolers (5 years of age; range: 58-74 months) participated in a randomized controlled trial in the summer before starting kindergarten (Fig 1). The recruitment period started on June 2^nd^ and ended on November 19^th^, 2019. An initial behavioral session ensured that all children participating in the study satisfied the following inclusion criteria: not knowing all uppercase letters and their corresponding sounds; having a Peabody Picture Vocabulary Test score higher than 85 (PPVT, 4th Edition; [22]); having normal or corrected to normal vision; being able to hold still for 5 minutes during an MRI mock scan. No neuropsychological or psychiatric disorder was reported. All children gave their assent to participate in the study, and their parents (or legal guardians) signed an informed consent form. The study was approved by the Institutional Review Board of the University of Washington.

**Fig 1.**
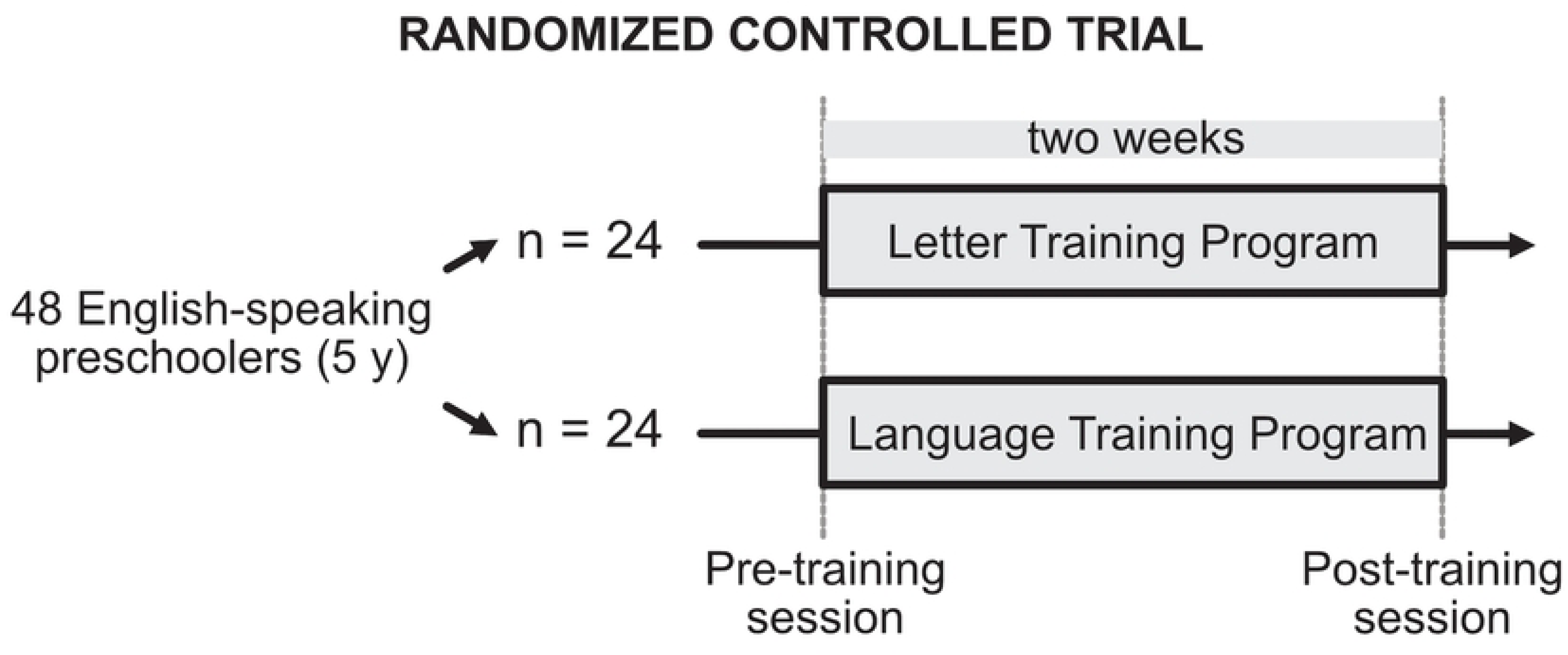
Graphical representation of the randomized controlled trial.

### Procedure

Participants were randomly assigned to one of two different training programs: a Letter (n = 24) or a Language (n = 24) organized into a fun and engaging summer camp. The Letter program followed the Slingerland method [23] and was focused on the foundational skills of reading such as letter recognition, letter-speech sound associations, and phonemic awareness (e.g., blending and segmentation of syllables and trigrams; Table 1). The Language program focused on oral linguistic abilities such as recognizing syntactic categories in spoken sentences, listening, comprehending and retelling stories, and learning new vocabulary (Table 1). Critically, the Language program did not include any exposure to written language compared to the Letter program which was almost exclusively focused on written language. Each training program was delivered to a small group of children (n = 6) by two teachers, who had a background in Education or Speech pathology. Each program lasted two weeks (3 hours/day, 5 days/week, 30 total hours) and was based on pedagogical models of Direct Instruction and Gradual Release of Responsibility. Letter and Language activities adopted a multisensory approach involving vision, audition and kinesthetics. Both programs had the same daily schedule and the learning process was scaffolded, so that the content of the activities followed an increasing degree of complexity. Each daily session started with 20 minutes of free play and ended with story time (Table 1). Starting from day 2, each daily activity started with a brief rehearsal of previous lessons.

**Table 2.**
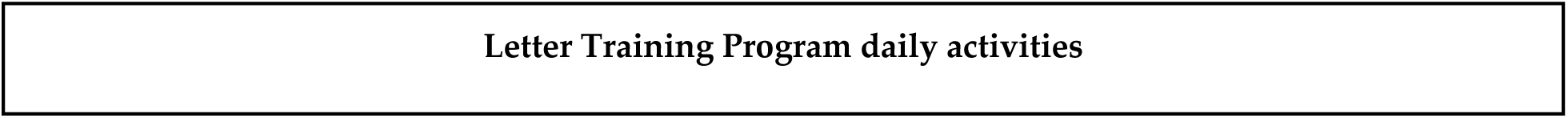
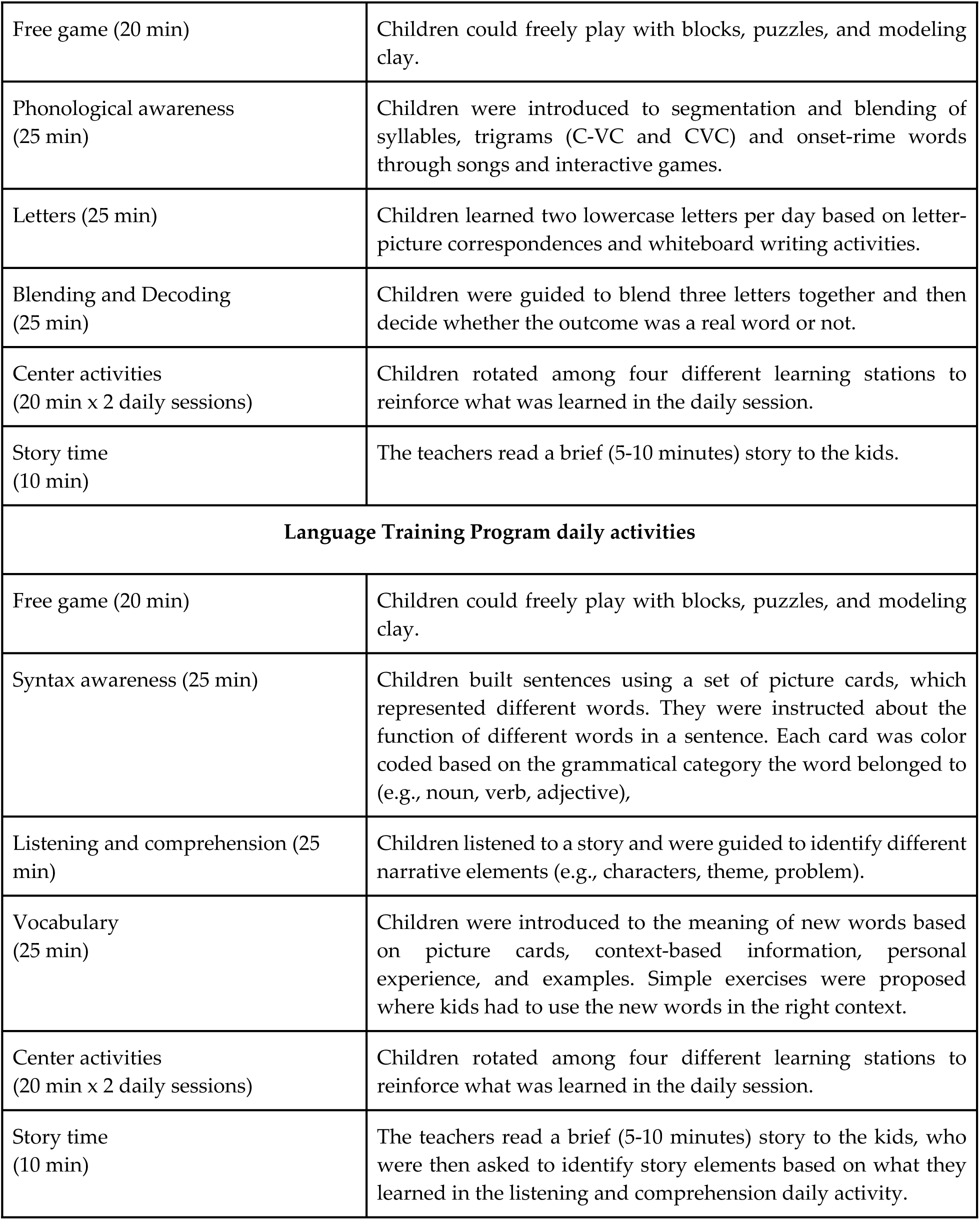
Description of the daily activities performed in the Letter and Language Training programs.

|  |
| --- |
| Letter Training Program daily activities |

|  |  |
| --- | --- |
| Free game (20 min) | Children could freely play with blocks, puzzles, and modeling clay. |
| Phonological awareness (25 min) | Children were introduced to segmentation and blending of syllables, trigrams (C-VC and CVC) and onset-rime words through songs and interactive games. |
| Letters (25 min) | Children learned two lowercase letters per day based on letter-picture correspondences and whiteboard writing activities. |
| Blending and Decoding (25 min) | Children were guided to blend three letters together and then decide whether the outcome was a real word or not. |
| Center activities (20 min x 2 daily sessions) | Children rotated among four different learning stations to reinforce what was learned in the daily session. |
| Story time (10 min) | The teachers read a brief (5-10 minutes) story to the kids. |
| **Language Training Program daily activities** |  |
| Free game (20 min) | Children could freely play with blocks, puzzles, and modeling clay. |
| Syntax awareness (25 min) | Children built sentences using a set of picture cards, which represented different words. They were instructed about the function of different words in a sentence. Each card was color coded based on the grammatical category the word belonged to (e.g., noun, verb, adjective), |
| Listening and comprehension (25 min) | Children listened to a story and were guided to identify different narrative elements (e.g., characters, theme, problem). |
| Vocabulary (25 min) | Children were introduced to the meaning of new words based on picture cards, context-based information, personal experience, and examples. Simple exercises were proposed where kids had to use the new words in the right context. |
| Center activities (20 min x 2 daily sessions) | Children rotated among four different learning stations to reinforce what was learned in the daily session. |
| Story time (10 min) | The teachers read a brief (5-10 minutes) story to the kids, who were then asked to identify story elements based on what they learned in the listening and comprehension daily activity. |

Behavioral and diffusion magnetic resonance imaging (dMRI) measures were collected before (average time: 19 days, SD: 9.23) and after (average time: 8 days, SD: 3.90) each training program.

### Behavioral data acquisition

During the behavioral session, the following standardized tests were administered: Phonological and Print Awareness Scale (tests of Initial sound matching, Final sound matching, and Phonemic awareness were administered from [24]); Phonological Awareness Literacy Screening (the Pseudoword decoding list was administered from [25]); Narrative Language Measures (the Story Retell test was administered from [26]); and the Expressive Vocabulary Test Third Edition [27]. Alphabet knowledge was also tested through flashcards presented in random order. There were 26 flashcards for uppercase letters, and 26 flashcards for lowercase letters. Each of the 26 cards was shown to the child and they were asked “What letter is this?” and “What sound does it make?”. The total score of the alphabet knowledge test was 52, both for upper and lowercase letters.

### dMRI data acquisition and preprocessing

MRI data was collected through a 3 T Phillips Achieva scanner with a 32-channel head coil (Philips, Eindhoven, Netherlands). A whole-brain anatomical volume at 1.0 x 1.0 x 1.0 mm resolution was acquired using a T1-weighted MPRAGE sequence (TR 9.2 s, TE 4.35 ms, matrix size 224 x 224, field of view 224 x 224 x 170, 170 slices). Diffusion-weighted magnetic resonance imaging (dMRI) data of the full brain were acquired with a spatial resolution of 2.0 mm^3^ (anterior-posterior phase encoding direction). A diffusion- weighted imaging (DWI) scan was acquired with 32 non-collinear directions (b-value = 1500 s/mm^2^; TR = 7200; TE = 83 ms). Four volumes with no diffusion weighting were also acquired (b-value = 0). To correct for echo-planar imaging distortions, one scan with a reversed phase encoding direction (posterior-anterior) and with three non-diffusion- weighted volumes was collected.

The T1-weighted (T1w) images were corrected for intensity non-uniformity (INU) using N4BiasFieldCorrection [28], ANTs 2.3.1), and used as T1w-reference throughout the workflow. The T1w-reference was then skull-stripped using antsBrainExtraction.sh (ANTs 2.3.1), using OASIS as target template. Spatial normalization to the ICBM 152 Nonlinear Asymmetrical template version 2009c [29] was performed through nonlinear registration with antsRegistration (ANTs 2.3.1, [30], using brain-extracted versions of both T1w volume and template. Brain tissue segmentation of cerebrospinal fluid (CSF), white-matter (WM) and gray-matter (GM) was performed on the brain-extracted T1w using FAST (FSL 6.0.3, [31]).

DMRI preprocessing and reconstruction were carried out using QSIprep 0.13.0RC2 ([32–34]), which is based on *Nipype* 1.6.0[32–34], *Nilearn* 0.7.1 [35] and *Dipy* 1.4.0 [36]. The preprocessing included topup distortion, MP-PCA denoising, motion and Eddy current correction (*q*-space smoothing factor = 10, 5 iterations; [37–40]). Only experimental sessions with a maximum framewise displacement below 4 mm and an average framewise displacement below 1 mm were further analyzed (Letter group pre-training session: 21; Letter group pre-training session: 22; Language group pre-training session: 19; Language group post-training session: 20). Multi-tissue fiber response functions were estimated using the dhollander algorithm as implemented in MRtrix3 [41]. Fiber orientation distributions (FODs) in each voxel were estimated via constrained spherical deconvolution (CSD,[42,43] using an unsupervised multi-tissue method[44,45]. Anatomically constrained tracking (ACT) was applied. FODs were intensity-normalized using mtnormalize[46]. Probabilistic tractography was carried out using the following QSIprep parameters: 1M streamlines, minimum length: 30 mm, maximum length: 250 mm. Fiber segmentation was carried out using pyAFQ 0.9 default parameters (https://yeatmanlab.github.io/pyAFQ;[47,48] cleaning iterations = 5, distance threshold = 5 SD, length threshold: 4 SD). Eighteen default tracts were segmented: Left/Right Arcuate, Left/Right Anterior Thalamic Radiation, Left/Right Cingulum, Left/Right Corticospinal Tract, Anterior/Posterior Forceps, Left/Right Inferior Fronto-Occipital Fasciculus, Left/Right Inferior Longitudinal Fasciculus, Left/Right Superior Longitudinal Fasciculus, Left/Right Uncinate. Diffusion metrics were calculated using the constrained spherical deconvolution model (CSD[49,50] and projected onto the tracts. Each streamline was resampled into a fixed number of nodes (n = 100), and average values of fractional anisotropy (FA), and mean diffusivity (MD) were calculated for each node. FA and MD were mapped onto each tract, weighting the values based on the streamline’s distance from the core of the tract [47]. Hence, FA and MD values of each white matter tract were calculated as the average across all 100 nodes of the tract profile.

## Results

### Behavioral results

A linear mixed effect model (LME) was run to test for behavioral effects due to the type of training received. Time (pre vs post session), Training Type (Letter vs Language) and their interaction were included as fixed effects. By-subject random intercepts and slopes were also included. Only models on alphabet knowledge (average accuracy of upper and lower case) and decoding skills showed a significant interaction between Training Type and Time (alphabet knowledge: β = 0.822, SE = 0.370, t = 2.224, p = 0.026; decoding skills:

β = 0.970, SE = 0.326, t = 2.972, p = 0.003) indicating that children participating in the Letter Training improved their letter knowledge (β = 2.770, SE = 0.784, t = 3.532, p < 0.001) and decoding ability (β = 1.689, SE = 0.496, t = 3.403, p = 0.001), while children participating in the Language Training group did not show such behavioral changes (alphabet knowledge: β = 0.896, SE = 0.776, t = 1.154, p = 0.248; decoding skills: β = 0.194, SE = 0.415, t = 0.469, p = 0.639; Fig 2).

**Fig 2.**
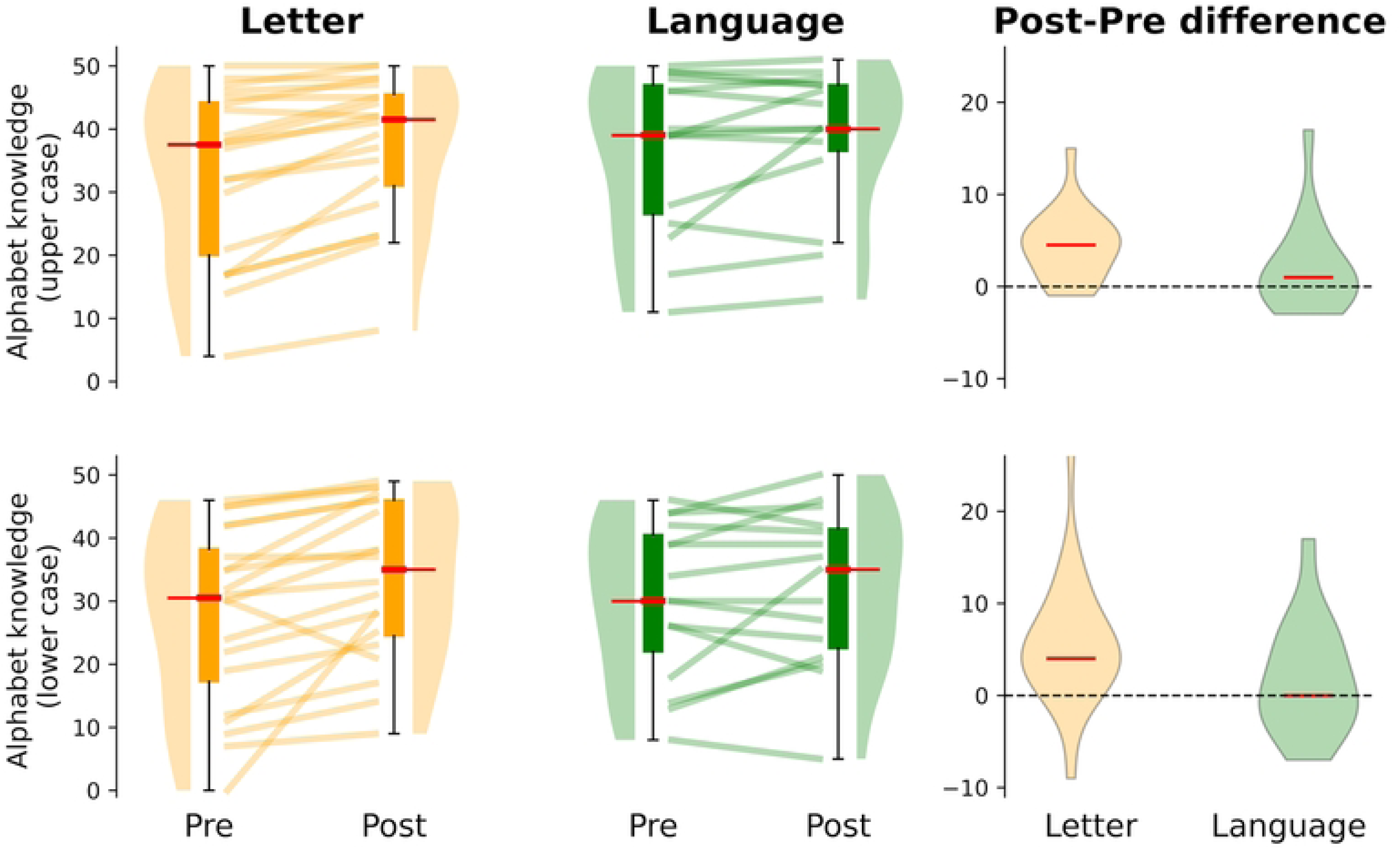
Training-related behavioral changes. First two columns: behavioral changes in alphabet knowledge from the Letter and Language Training groups for each experimental session. The third column shows the distribution of alphabet knowledge changes (i.e. difference between the individual scores obtained in the post and pre-training sessions) for each group.

### dMRI results

#### Scan Rescan reliability

For each white matter tract and diffusion property (FA and MD), we calculated scan-rescan reliability metrics to quantify the consistency of two types of dMRI measures between experimental sessions: profile and subject reliability (as in [51]). For the profile reliability, a Pearson correlation was calculated between the pre and post session tract profiles of each participant. To calculate the subject reliability, a Pearson correlation was calculated between the pre and post sessions median values of the tract profiles of each participant. Individual correlation coefficients were averaged for each tract to obtain a reliability estimate. Our dMRI measures showed high degrees of scan-rescan reliability between the two experimental sessions (profile reliability: FA, median *r* = 0.99, range 0.93- 0.99; MD, median *r* = 0.92, range: 0.65-0.99; subject reliability: FA, median *r* = 0.83, range 0.62-0.90; MD, median *r* = 0.87, range: 0.55-0.92; Fig 3, for Subject reliability of each experimental group, see S1 and S2 Figs).

**Fig 3.**
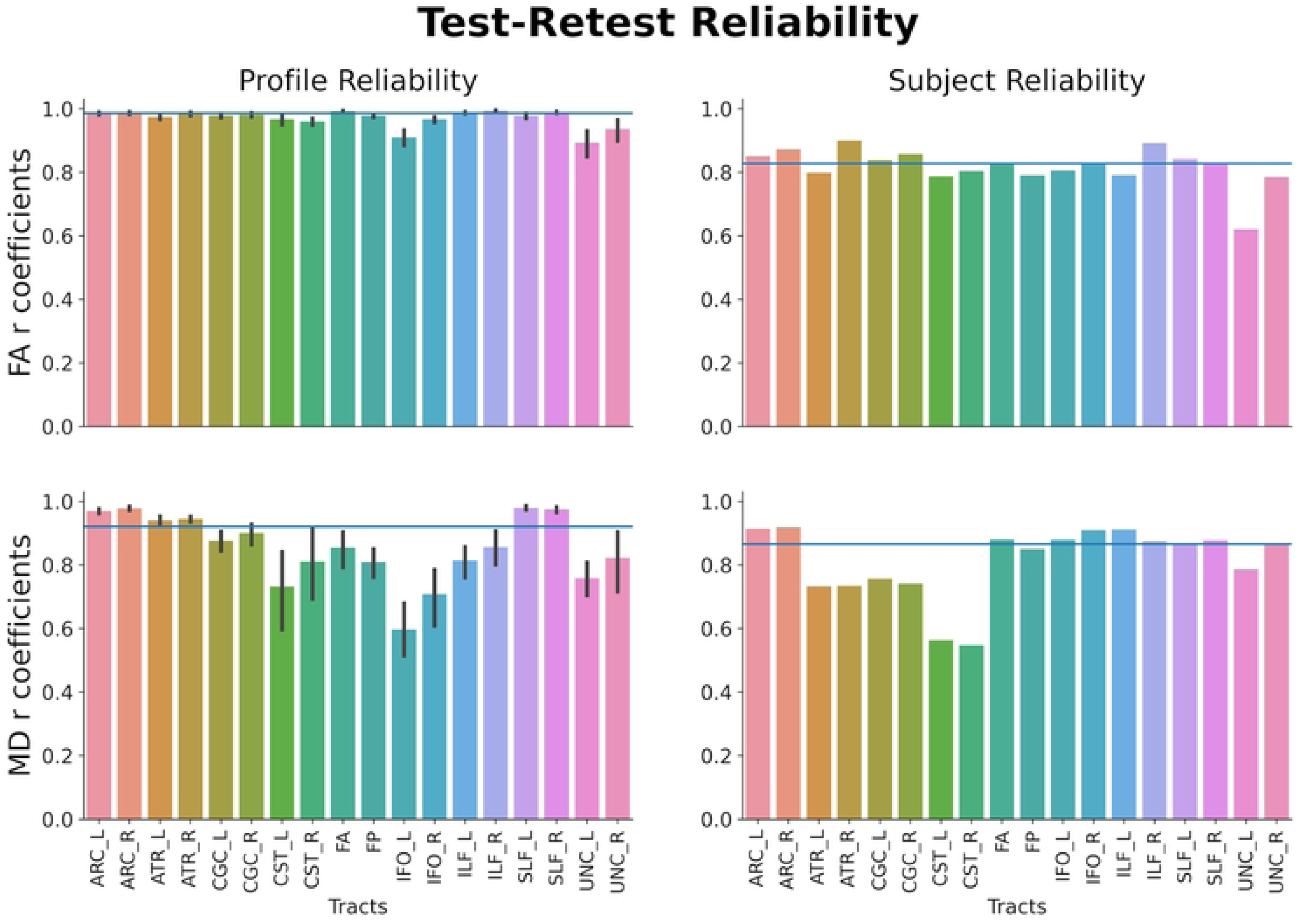
Scan Rescan reliability. The two columns show the profile and subject reliability estimates of each white matter tract examined in the study. The two rows show reliability estimates for FA and MD, respectively. ARC: Arcuate Fasciculus; ATR: Anterior Thalamic Radiation; CGC: Cingulum Cingulate; CST: Corticospinal Tract; FA: Anterior Forceps; FP: Posterior Forceps; IFO: Inferior Longitudinal Fasciculus; ILF: Inferior Longitudinal Fasciculus; SLF: Superior Longitudinal Fasciculus; UNC: Uncinate.

#### Training effects on dMRI measures

An LME model was run on the average FA and MD values of each tract profile to test for structural changes due to the type of training received. Time (pre vs post session), Training Type (Letter vs Language) and their interaction were included as fixed factors. By-subject random intercepts were also included. No structural changes were observed between experimental sessions (FA: all *t*s<2; MD: all *t*s<2.5) or between the two groups (FA: all *t*s<2.8; MD: all *t*s<2). Interactions between Training Type and Time were not significant (FA: all *t*s<2.12; MD: all *t*s<2), suggesting that no statistically significant changes were observed for either group over the 2-week training period (Fig 4 shows the results for the left arcuate; for similar results on the left ILF, see S3 Fig).

**Fig 4.**
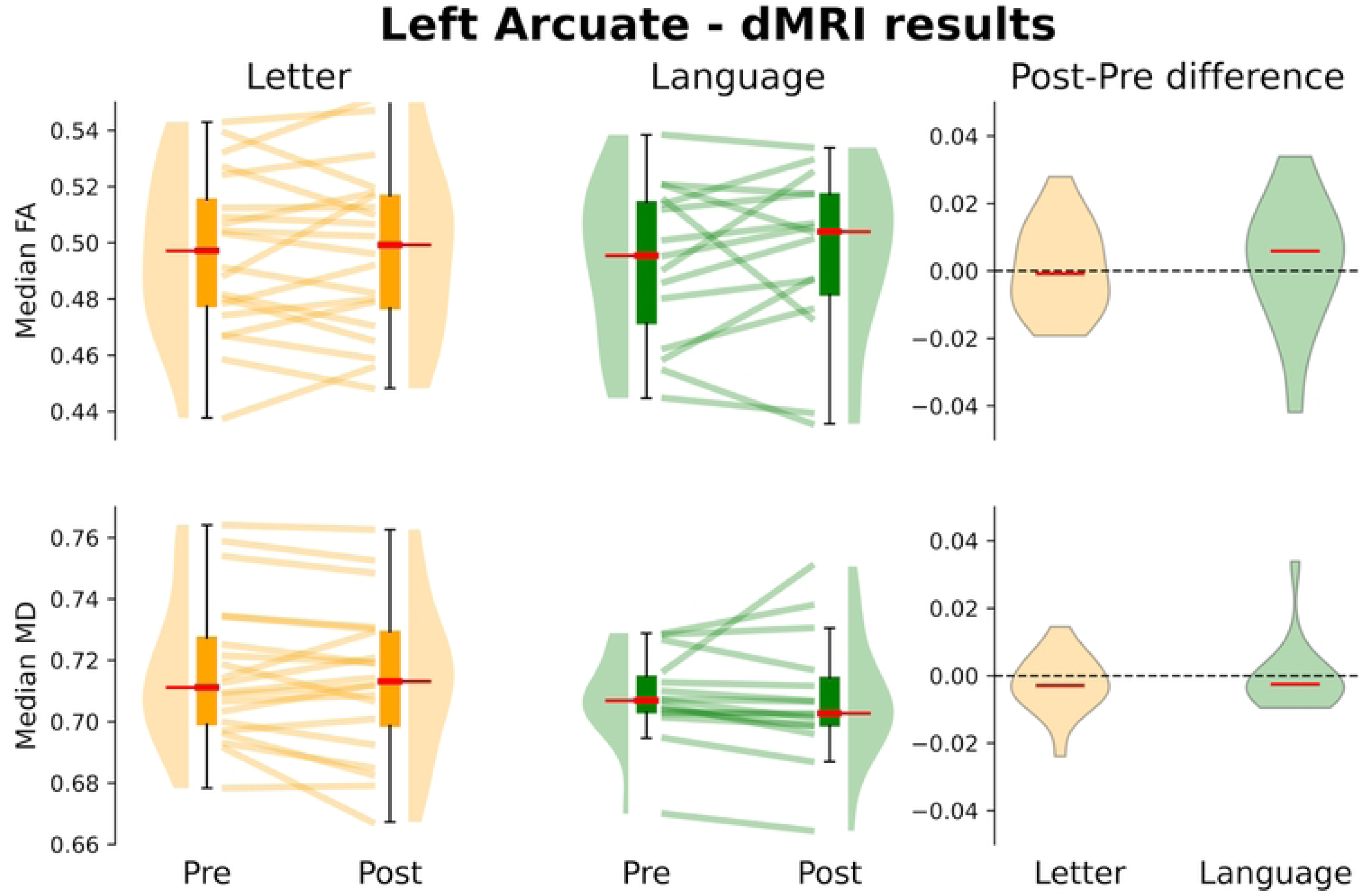
Training-related dMRI changes of the left arcuate. First two columns: structural changes of the left arcuate are shown for the Letter and Language Training groups and each experimental session. The third column shows the distribution of FA and MD changes (i.e. difference between the individual profiles observed in the post and pre-training sessions) for each group.

Similar LME models were fitted for each single node of each tract profile and they confirmed this pattern of results (Figs 5 and 6); there were no observable changes in white matter properties over the two-week training period.

**Fig 5.**
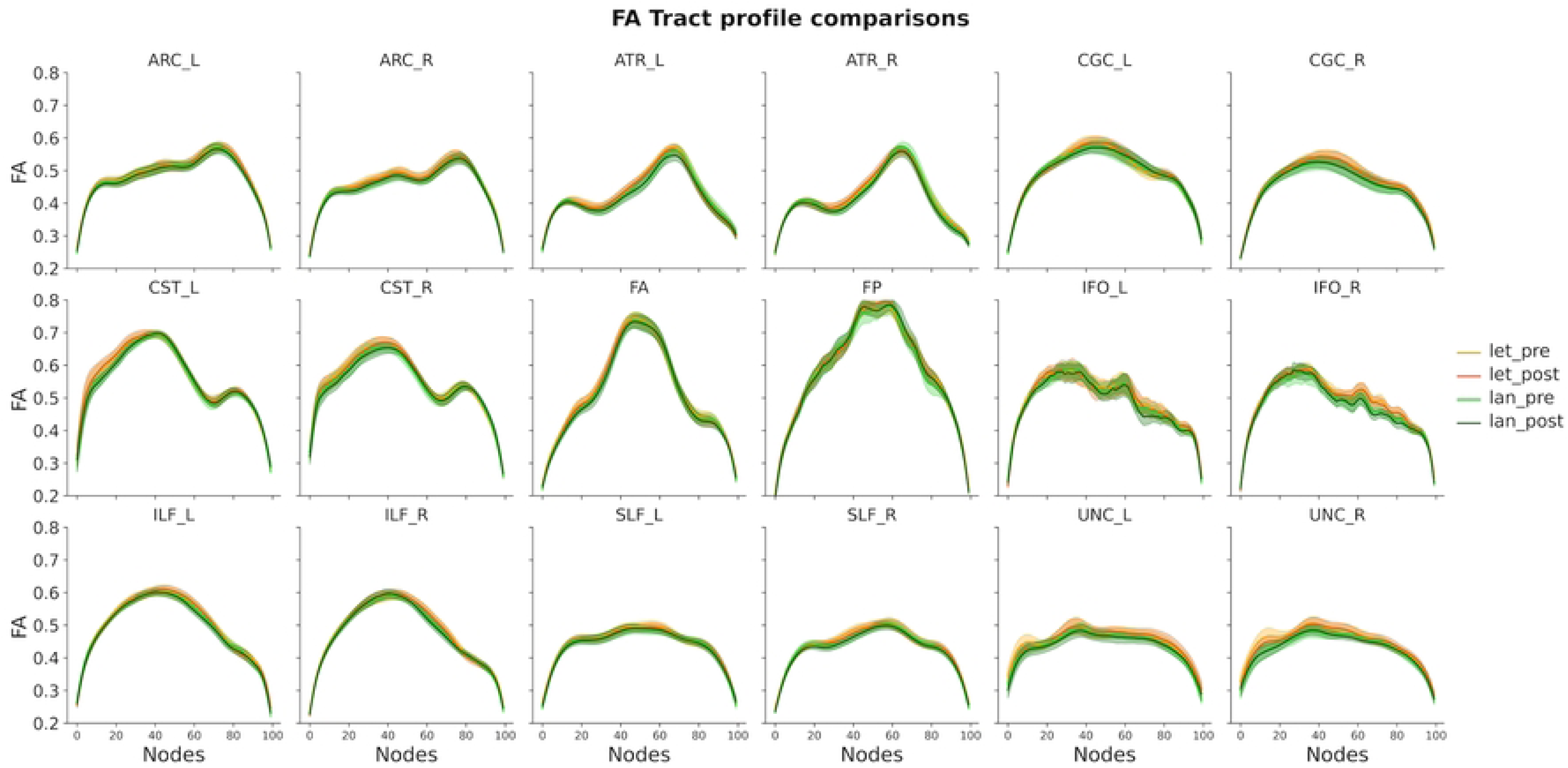
FA tract profile for each experimental group and training session. The plots show FA values estimated based on the beta coefficients extracted from node-by-node LME models. Shaded areas represent +/- 2 SE. ARC: Arcuate Fasciculus; ATR: Anterior Thalamic Radiation; CGC: Cingulum Cingulate; CST: Corticospinal Tract; FA: Anterior Forceps; FP: Posterior Forceps; IFO: Inferior Longitudinal Fasciculus; ILF: Inferior Longitudinal Fasciculus; SLF: Superior Longitudinal Fasciculus; UNC: Uncinate.

**Fig 6.**
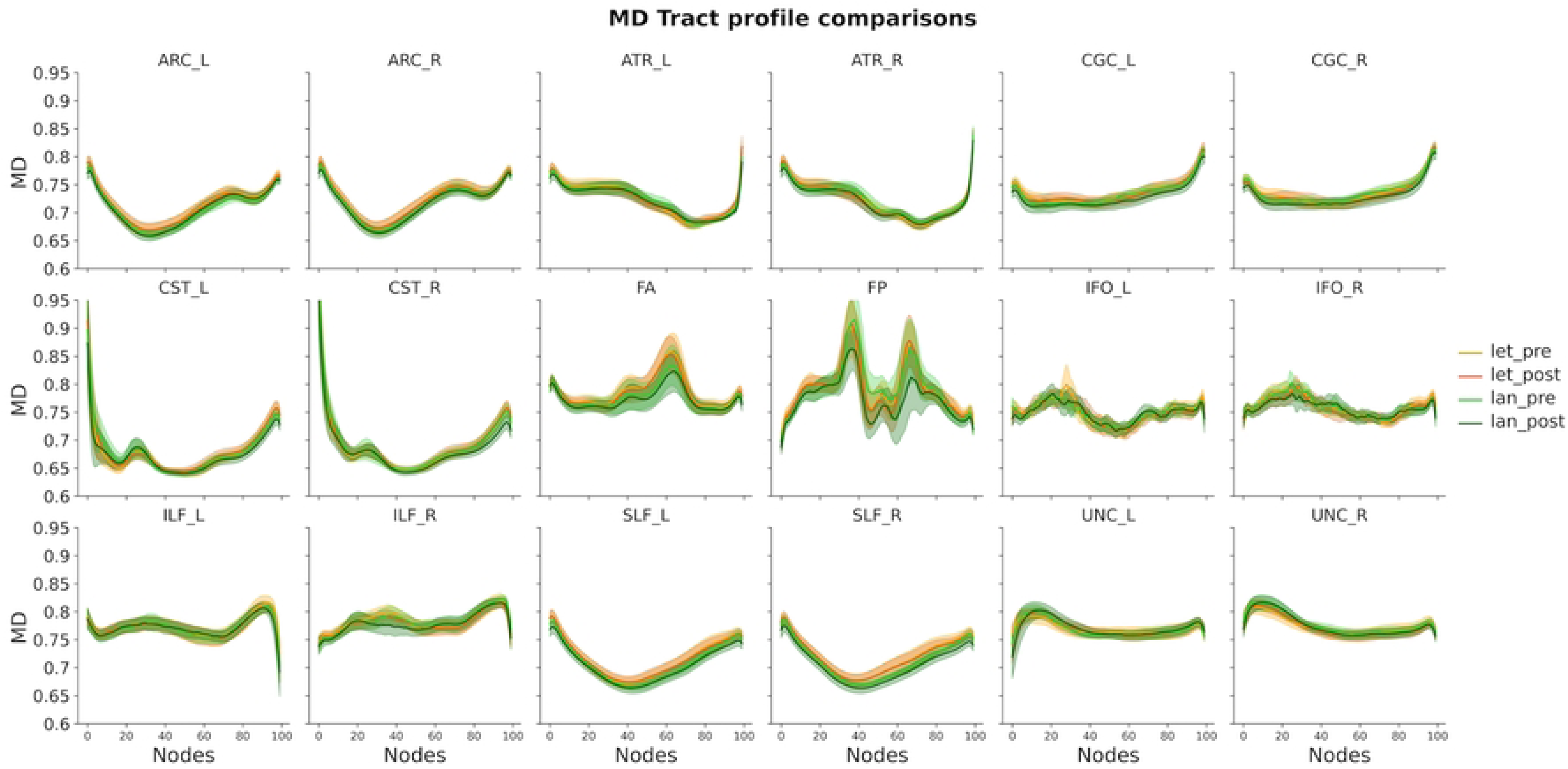
MD tract profile for each experimental group and training session. The plots show MD values estimated based on the beta coefficients extracted from node- by-node LME models. Shaded areas represent +/- 2 SE. ARC: Arcuate Fasciculus; ATR: Anterior Thalamic Radiation; CGC: Cingulum Cingulate; CST: Corticospinal Tract; FA: Anterior Forceps; FP: Posterior Forceps; IFO: Inferior Longitudinal Fasciculus; ILF: Inferior Longitudinal Fasciculus; SLF: Superior Longitudinal Fasciculus; UNC: Uncinate.

To estimate the evidence supporting the null hypothesis (H0: no plasticity), additional Bayesian analyses were run on each tract to compare the dMRI training effect (post-pre mean profile difference) between the two groups. Bayes factors supported small-to- moderate evidence for the null effect in the majority of the tracts, including all tracts that are part of the reading circuitry (FA: BFs<1 in 16 of the 18 tracts; MD: BFs<1 in 12 of the 18 tracts; Fig 7).

**Fig 7.**
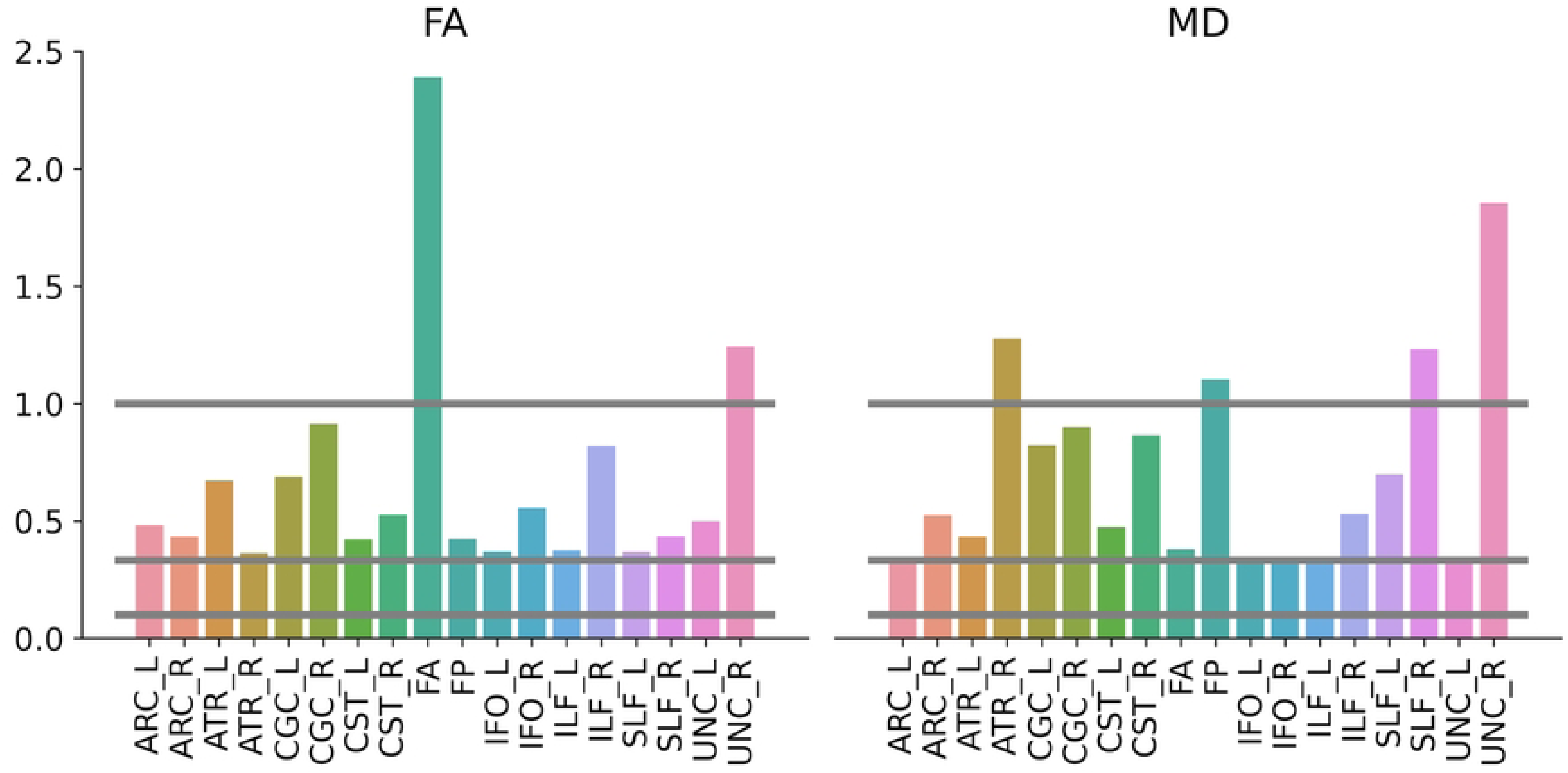
Bayes factors relative to the group comparison of the dMRI training effect for each tract. ARC: Arcuate Fasciculus; ATR: Anterior Thalamic Radiation; CGC: Cingulum Cingulate; CST: Corticospinal Tract; FA: Anterior Forceps; FP: Posterior Forceps; IFO: Inferior Longitudinal Fasciculus; ILF: Inferior Longitudinal Fasciculus; SLF: Superior Longitudinal Fasciculus; UNC: Uncinate.

### Linking training effects between behavioral and dMRI measures

Despite the lack of experience-driven white matter plasticity at the group level, it is possible that longitudinal change in alphabet knowledge might relate to longitudinal white matter changes. To test this hypothesis, the two groups were combined and Pearson correlations were calculated to check whether individual changes in alphabet knowledge (average of lower and upper case knowledge) could be mapped onto structural changes of left AF and ILF. This analysis did not show significant effects after Bonferroni correction (see Table 3).

**Table 3.** Pearson correlations between post-pre differences in reading performance and structural properties.

|  | FA |  | MD |  |
| --- | --- | --- | --- | --- |
|  | *r* | *p corrected* | *r* | *p corrected* |
| Left ARC | -0.38 | 0.104 | 0.12 | 0.99 |
| Left ILF | -0.41 | 0.060 | 0.36 | 0.124 |

## Discussion

This randomized controlled trial examined short-term effects of a Letter and a Language training program on preschoolers’ reading performance and brain structure. The findings suggest that a two-week Letter training program causes improvements in preschoolers’ letter knowledge and decoding skills. However, this behavioral effect was not accompanied by short-term changes in the diffusion properties (i.e., FA and MD) of white matter pathways, within or outside the reading circuitry. The presence of quick behavioral changes as a result of Letter training confirms previous findings on the effectiveness of short-term reading instruction, which has been observed in children with and without reading disorders [52–55].

Our dMRI findings further complement the existing literature on short-term reading- relating brain plasticity by showing that reading performance improvements are not always accompanied by changes in diffusion properties of white matter pathways [4,5,12,20,21]. The high reliability estimates for both FA and MD scores across sessions ensure that this null effect could not be accounted for by low dMRI data quality. Bayesian analyses provided support for the null hypothesis (no change in white matter diffusion) for all major white matter tracts of the reading network. In addition, correlation analyses confirmed the lack of a clear correspondence between preschoolers’ individual behavioral changes and variations in structural properties of reading white matter pathways.

One aspect that can account for the lack of structural changes is the type and intensity of the reading program. Previous studies have mainly focused on effects of reading intervention in children diagnosed with dyslexia, which can have an intense and profound impact on struggling readers’ cognitive and social lives. In the current study, our reading training proposed preschool/kindergarten activities that are usually carried out in a classroom setting. Since these training programs are similar to common preschool and kindergarten classrooms, they might not represent a dramatic enough environmental change to cause large-scale remodeling of the white matter. Related to this point, another variable that can account for our results is the cognitive profile of the trainees. This is the first randomized controlled trial on the effects of a short-term reading training with typically developing preschoolers. Previous experimental evidence collected so far (Table 1) refers to the effects of short-term remediation programs on children with reading disorders or at-risk of reading disorders. Hence, the large effects that have been reported so far might reflect the dramatic environmental change of entering an intensive intervention environment after struggling in school for years. This experience is quite different than typically developing children beginning formal reading instruction [3–5,19].

Another possible explanation to consider is the type of diffusion properties examined here. Recent dMRI findings on the short-term effects of reading intervention programs in preschoolers reported structural changes only in myelin water fraction, but not in FA and MD scores [3,21]. This might suggest that MRI measures more specifically related to myelination would better reflect reading-related short-term plasticity around 5 years of age. However, within 7 and 12 years of age an opposite pattern of results have been reported, with MD and FA providing evidence for rapid structural plasticity while no training-dependent changes were reported for more myelin-specific correlates, such as axonal water fraction and R1 [4,19]. These results are still compatible with the idea that short-term plasticity due to reading training might affect different structural properties of white matter depending on the developmental time window (e.g., there might be a higher degree of plasticity for myelin-specific indexes in the early stages of life). Additional research is needed in order to clarify which type of diffusion properties can be shaped by experience as a function of age (e.g., neural or non-neuronal plasticity, intra or extra axonal plasticity, [3,19,56]).

Finally, another potential explanation for our dMRI findings regards the presence of a possible time shift between the training effects on behavior and brain structure, with white matter changes happening over a larger temporal scale compared to behavioral changes. For instance, our findings are still compatible with the idea that at this early age the amount of training received is not sufficient to shape what will become the reading circuit later on. Although some studies have shown no time lag between behavioral and structural changes in response to a short reading intervention program [3–5,12], the exact time course of reading-related neuroplasticity is still understudied and needs further investigation.

Overall, this heterogeneous picture of findings on short-term reading-related structural neuroplasticity highlights the need to better define the conditions under which white matter can be shaped by experience. Several experiential and developmental factors might modulate the degree of white matter plasticity exhibited in response to reading training or intervention. Research evidence coming from other cognitive domains might give us some insights on the critical constraining variables to be considered. For instance, studies testing for the presence of a sensitive period of sensory and motor white matter circuits suggest that the time onset of the environmental exposure is a key factor to establish whether white matter structure is stable or plastic [57,58]. Other factors that have been suggested to modulate the balance between structural plasticity and stability are the type and the duration of experiential exposure [59–63] and the individual cognitive health and lifestyle risk factors [61,64].

In conclusion, this randomized controlled trial highlights that a two-week literacy training can cause fast behavioral changes in preschoolers’ reading performance, without being accompanied by fast FA and MD changes of the reading circuitry. These findings highlight that rapid diffusion properties variations are not always observed in response to short-term reading training and point to the need of specifying the conditions under which white matter structure is plastic versus stable.

## Supporting information

**S1 Fig. Subject reliability of FA for each experimental group.**

**S2 Fig. Subject reliability of MD for each experimental group.**

**S3 Fig. Training-related dMRI changes of the left inferior fasciculus.**

First two columns: structural changes of the left ILF are shown for the Letter and Language Training groups and each experimental session. The third column shows the distribution of FA and MD changes (i.e. difference between the individual profiles observed in the post and pre-training sessions) for each group.

